# Optimizing *ex-vitro* one-step RUBY-equipped hairy root transformation in drug- and hemp-type Cannabis

**DOI:** 10.1101/2023.11.29.569008

**Authors:** Ladan Ajdanian, Mohsen Niazian, Davoud Torkamaneh

## Abstract

Using synthetic biology techniques to engineer secondary metabolic pathways through hairy root transformation is one of the most advanced approaches used in research. In this study, we optimized an *ex-vitro* one-step hairy root transformation of the *RUBY* system in both drug- and hemp-type cannabis, shedding light on its potential applications in secondary metabolite production. Three different strains of *A. rhizogenes* including (A4, ARqual, and K599) were used. Significant variation in HR induction and transformation efficiency (TE) was observed based on *A. rhizogenes* strains and seed types. Drug-type seedlings exhibited the highest hairy root induction, increasing by 58.8% compared to hemp-type seedlings. Also, the A4 strain consistently demonstrated the highest transformation efficiency (75%) irrespective of genotype, while the ARqual strain yielded the lowest one (8.33%). In conclusion, our study is the first to present an *ex-vitro* one-step transformation of both hemp- and drug-type cannabis. In comparison to the in vitro method, our ex-vitro method is simpler, faster, and has a lower risk of contamination, making it an excellent choice for the efficient production of secondary metabolites in cannabis using the CRISPR/Cas system.

## Introduction

Cannabis (*Cannabis sativa* L.), once concealed by the veil of prohibition, is now emerging as a versatile and promising plant species, riding the wave of recent legalization. This transformation has unlocked unprecedented opportunities for both medical research and industry growth, positioning cannabis on a trajectory to reach a projected market size of USD 444.34 Billion by 2030 (*fortune business insights*). Despite the plant’s capability to produce more than 545 potentially bioactive secondary metabolites (Walsh et al., 2021), its legal categorization in Canada, the USA, and Europe hinges on the concentration of a solitary cannabinoid, Δ^9^-tetrahydrocannabinol (THC), found in female flowers (Torkamaneh & Jones, 2022). Categorically, cannabis plants containing less than 0.3% THC are classified as hemp-type (or industrial or fiber type), while those exceeding 0.3% THC are labeled as drug-type (or medicinal or recreational) cannabis. Beyond the major cannabinoids (THC and cannabidiol (CBD)), cannabis synthesizes over 120 additional cannabinoids referred to as minor and/or rare cannabinoids. These compounds, produced in smaller quantities, alongside THC and CBD, from the parent cannabinoid cannabigerolic acid (CBGA), remain poorly understood due to their scarcity.

In recent decades, hairy root (HR) culture, an established method facilitated by *Agrobacterium rhizogenes*-mediated transformation techniques, has gained significant attention by academic research teams, biotechnology companies and pharmaceutical industries as a convenient and viable approach to produce target metabolites due to its rapid growth and stability in terms of both biochemistry and genetics (Faraz et al., 2020). In recent years, exploration into the production of secondary metabolites in hemp-type cannabis through HR culture, conducted in *in vitro* conditions, has shown promising results (Gutierrez-Valdes et al., 2020). The challenge in inducing HR through *in vitro* culture, characterized by its time-consuming and labor-intensive nature, along with concerns regarding poor productivity and scalability, has historically limited the widespread adoption of HR cultures. Obtaining suitable HRs from *A. rhizogenes* represents the pivotal the initial step in producing secondary metabolites from transformed HRs. While a method for the *in-vitro* HR induction of cannabis has been proposed (Farag & Kayser, 2015), there is currently no reported *ex-vitro* method.

## Material and Method

Here, we optimized an *ex-vitro* one-step HR transformation of the *RUBY* system in both drug- and hemp-type cannabis, shedding light on its potential applications in secondary metabolite production. Freshly produced hemp-type (cv. Vega; 0.2% THC and 1.6% CBD) and drug-type (cv. AD248; 24% THC and 0.05% CBD) cannabis seeds from Université Laval in 2023 (Lapierre et al., 2023) were germinated in seed starter peat pellets (Jiffy group, Canada). Ten-day-old seedlings were diagonally excised from the apical portion of the hypocotyl using a sterile scalpel and subsequently inoculated (Figure S1a) with three different *A. rhizogenes* strains (A4, ARqual, and K599). In this study, we used a *RUBY* binary vector carrying the visually detectable *RUBY* reporter gene and the *bar* selectable marker (RBV; pARSCL504 [pTRANS_230] 35S:Ω:Ruby (Addgene # 198636)). The preparation and transformation of bacteria followed a previously optimized protocol by our group (Niazian et al., 2023). *A. rhizogenes* inoculation was conducted using a one-step *ex vitro* method established in our lab (Niazian et al. (unpublished); Method S1). Inoculated seedlings were enclosed in paper bags (Figure S1b) and placed in a growth chamber (26:22°C day: night, 16:08 light: dark, and 80% of humidity). Plants were irrigated every other day with nutrient solution (Table S1) and sprayed with sterilized distilled water.

## Result and Discussion

HRs emerged 10 days post-inoculation, at the cutting place, and putative transgenic HRs (characterized by the expression of the *RUBY* gene, observed as red HRs) appeared 14 days post-inoculation (Figure 1a_1_, Figure S1c). Mature putative transgenic HRs were observed on the 20^th^ day of the experiment (Figure 1a_2_). The plants were maintained under these conditions for an additional 10 days, resulting in composite plantlets with a substantial mass of red HRs (Figure 1b, Figure S1d). The percentage of transformed HR and transformation efficiency were evaluated under various conditions, including genotype (hemp-vs. drug-type) and bacterial strain. Finally, the putative transgenic HRs was evaluated for the presence of the T-DNA by PCR amplifying the *bar* gene (Methods S1 & Figure S1e).

**Figure 1.**
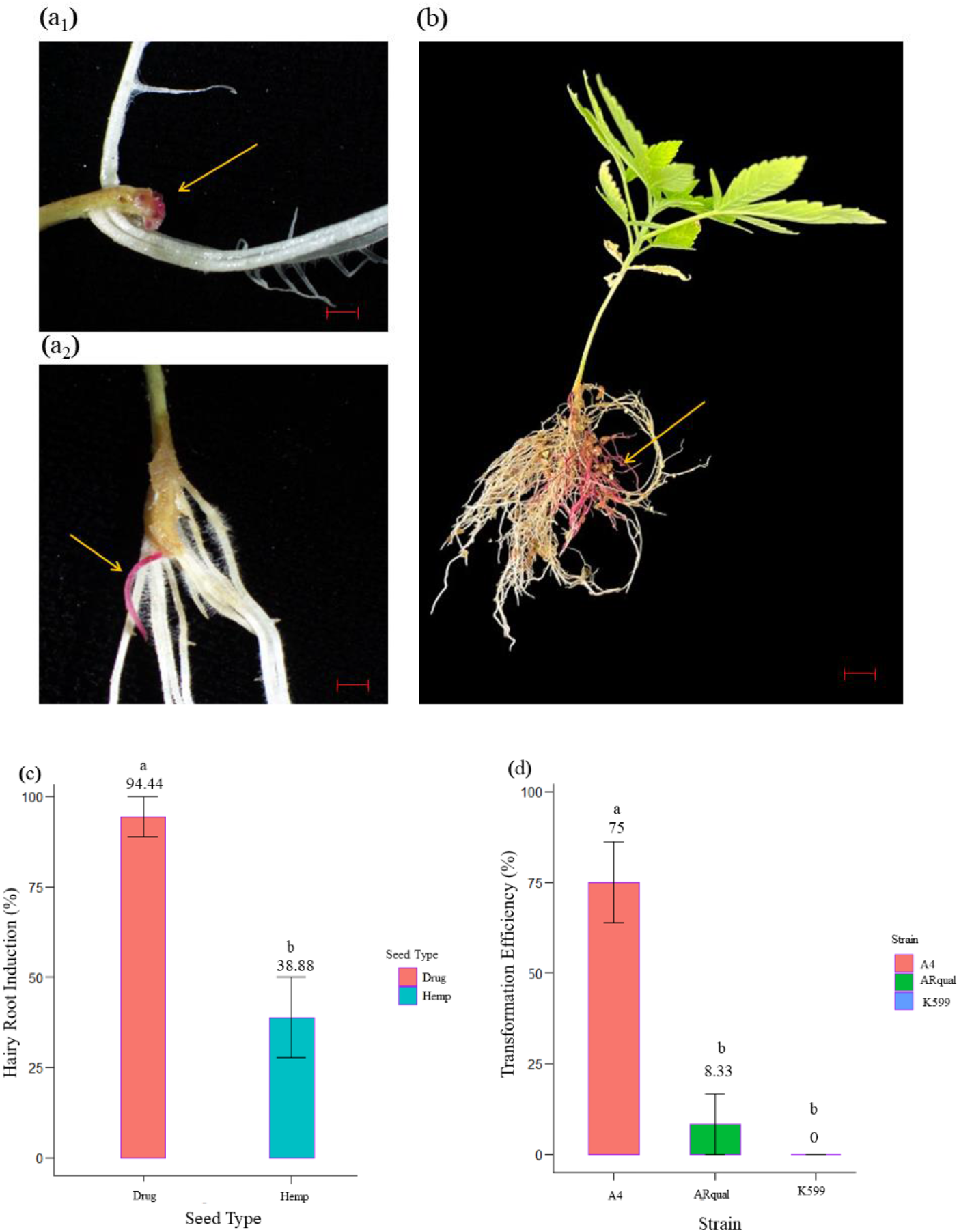
One-step hairy root induction and *RUBY* gene expression in cannabis. (a_1_) The emergence of red hairy roots expressing *RUBY* gene in cannabis (bar= 1 cm). (a_2_) The elongation of red hairy root expressing *RUBY* reporter gene (bar= 1 cm). (b) A fully developed composite cannabis plant (bar = 1 cm). (C) Mean comparison ± standard error (LSD P ≤ 0.05) with two different types of seed on percentage of explants showing hairy root induction in cannabis. (D) Mean comparison ± standard error (LSD P ≤ 0.05) with three different stains on Percent of transformed roots in cannabis. Different letters denote treatments that produce statistically different means.

Significant variation in HR induction and transformation efficiency (TE) was observed based on *A. rhizogenes* strains and seed types (Table S2). All strains used in this study were led to HR induction, with no significant differences observed. However, the plant type (hemp vs. drug type) significantly influenced (*p* = 0.0003) the percentage of HR induction (Figure 1d). Overall, drug-type seedlings exhibited the highest HR induction, increasing by 58.8% compared to hemp-type seedlings. The induction of HRs in *in-vitro* condition has been previously reported, showing a noticeable difference in the percentage of induction for hemp-type cannabis plants (Wahby et al., 2017).

The A4 strain consistently demonstrated the highest TE (75%) irrespective of genotype, while the ARqual strain yielded the lowest TE (8.33%). Notably, the K599 treatment did not result in the formation of transformed roots (Figure 1c). Yet, no significant difference (*p* = 1) was observed between hemp- and drug-type cannabis (Figure 1d). This is important, as both type of plants can be efficiently transformed using the best strain (A4) found in this study.

In conclusion, even though *in vitro* HR transformation of hemp-type cannabis using *A. rhizogenes* (strain A4) has been documented previously (Berahmand et al., 2016), our study presents the first *ex vitro* one-step transformation in both hemp- and drug-type cannabis. Compared to the *in vitro* method, our *ex-vitro* method offers simplicity, speed, and reduced contamination risk, making it an optimal choice for the efficient production of secondary metabolites using CRISPR/Cas system in cannabis.

## Acknowledgements

We thank NSERC for financial support.

## Conflicts of interest

The authors declare no conflict of interest.

## Supporting information

**Methods S1**. Binary vector construct and A. *rhizogenes* transformation.

**Table S1**. Nutritional program used in this experiment.

**Table S2**. Analysis of variance (ANOVA) of hairy root induction and transformation efficiency.

**Figure S1**. (a) Cannabis seedling infected with the specific bacteria, and (b) placed seedling in paper bags. (c) First sign of red (transformed) root. (d) A fully grown transformed hairy roots. (e) PCR results to validate transformation. M stands for a 100 bp DNA marker; 1–7 denotes red transgenic hairy roots; P is the 35S_Ω_RUBY plasmid used as a positive control; C stands for white hairy roots used as a control sample; and H_2_O is a negative control containing no template.

## Notes

### Competing Interest Statement

The authors have declared no competing interest.

