## Supporting Information for "Optimizing *ex-vitro* one-step RUBY-equipped hairy root transformation in drug- and hemp-type Cannabis"

Département de phytologie, Université Laval, Québec City, QC, Canada, Institut de Biologie Intégrative et des Systèmes (IBIS), Université Laval, Québec City, QC, Canada, Centre de recherche et d'innovation sur les végétaux (CRIV), Université Laval, Québec City, QC, Canada, Institute Intelligence and Data (IID), Université Laval, Québec City, QC, Canada

### Supporting information

#### Methods S1. Binary vector construct and *A. rhizogenes* transformation.

The plasmid used in this study was a binary construct (pARSCL504 [pTRANS_230] 35S:W:Ruby, Addgene #198636) carrying the reporter gene *RUBY* along with the *bar* selectable marker in its T-DNA region (Niazian et al. 2023). The *RUBY* reporter gene allows real-time monitoring without any disruption; thus, it is referred to as the Ruby Binary Vector (RBV). The RBV was used to transform *A. rhizogenes* competent cells that had been prepared using a freeze-thaw technique (An et al. 1989). On YEP plates containing 50 mg/L kanamycin, transformed colonies were selected. Glycerol stocks were made by suspending fresh single colonies in a liquid YEP medium containing 50% (vol/vol) glycerol and storing the mixture at -80°C.

***Ex vitro* hairy root transformation.** A single colony (an *Agrobacterium* carrying a binary RUBY vector) from each strain was taken and suspended in 850 µL of liquid YEP medium containing 15% (vol/vol) glycerol. 300 µL of the final mixture was then applied to YEP solid plates that had 50 mg/L of kanamycin. The next step was an overnight incubation at 28°C. Seedlings of cannabis were then infected using a dense bacterial lawn. Each seedling was infected with the specific bacteria before being placed in designated growth sheets. Plants were then placed in a controlled environment, and every day the presence of red and hairy roots was monitored. Red roots were visually observed 20 days after the experiment started. Next, using formulas 1 and 2, the hairy root induction percentage and efficiency were determined.

**(1)** $$\text{HR}\ (\%) = \frac{\boldsymbol{N_{hr}}}{\boldsymbol{N_T}} \times \mathbf{100}$$

HR is the percentage of hairy root induction; $N_{hr}$ is the number of seedlings with hairy roots (lengths ≥ 1 cm); $N_T$ is the total number of inoculate seedlings.

**(2)** $$\text{TE}\ (\%) = \frac{N_{RUBY}}{N_I} \times 100$$

TE is transformation efficiency; $N_{RUBY}$ is the number of seedlings with at least one hairy root expressing the

RUBY gene; $N_I$ is the number of inoculated seedlings with hairy roots (length ≥ 1 cm) (Su et al. 2022; Melito et al. 2010)

Finally, a PCR validation experiment was performed using primers amplifying (413-bp) the *bar* gene (F:5'GACAAGCACGGTCAACTTCC-3'; R: 5' AGTCCAGCTGCCAGAAACC-3') from genomic DNA extracted from the red roots and white roots (control) to determine whether the T-DNA was present in any red roots. Analysis of variance (ANOVA) and means comparison analysis were carried out using SAS® (SAS Institute Inc., Cary, NC). The normal distribution of data was checked before the analysis of variance. Means were compared using LSD test ($P \leq 0.05$). All graphs were generated using R software (version 4.3.2).


An G, Ebert PR, Mitra A, Ha SB (1989) Binary vectors. Plant molecular biology manual:29-47

Chamness JC, Kumar J, Cruz AJ, Rhuby E, Holum MJ, Cody JP, Tibebu R, Gamo ME, Starker CG, Zhang F (2023) An extensible vector toolkit and parts library for advanced engineering of plant genomes. The Plant Genome:e20312

Melito S, Heuberger AL, Cook D, Diers BW, MacGuidwin AE, Bent AF (2010) A nematode demographics assay in transgenic roots reveals no significant impacts of the Rhg1 locus LRR-Kinase on soybean cyst nematode resistance. BMC plant biology 10:1-14

Niazian M, Belzile F, Curtin SJ, de Ronne M, Torkamaneh D (2023) Optimization of in vitro and ex vitro Agrobacterium rhizogenes-mediated hairy root transformation of soybean for visual screening of transformants using RUBY. Frontiers in Plant Science 14

Su Y, Lin C, Zhang J, Hu B, Wang J, Li J, Wang S, Liu R, Li X, Song Z (2022) One-step regeneration of hairy roots to induce high tanshinone plants in Salvia miltiorrhiza. Frontiers in Plant Science 13:913985

**Table S1.** Nutritional program used in this experiment.

| Fertilizer | 12-2-14 | 15-30-15 | Solupotasse | Micro | Activ 0-0-5 | EZ-Gro Armour 0-0-15 |
|---|---|---|---|---|---|---|
| Per 1 Lit | 0.7gr | 0.2gr | 0 | 0.1gr | 3gr | 2cc |

**Table S2.** Analysis of variance (ANOVA) of hairy root induction and transformation efficiency.

| Source of variation | df | Mean squares | |
|---|---|---|---|
| | | Hairy root induction | Transformation efficiency |
| Strain | 2 | $1666.67^{ns}$ | $10138.89^{**}$ |
| Seed type | 1 | $13888.89^{**}$ | $0^{ns}$ |
| Strain × Seed type | 2 | $555.56^{ns}$ | $416.67^{ns}$ |
| Error | 12 | 555.56 | 416.67 |

*Significant at p≤0.05. **: Significant at p≤0.01. ns: Insignificant.

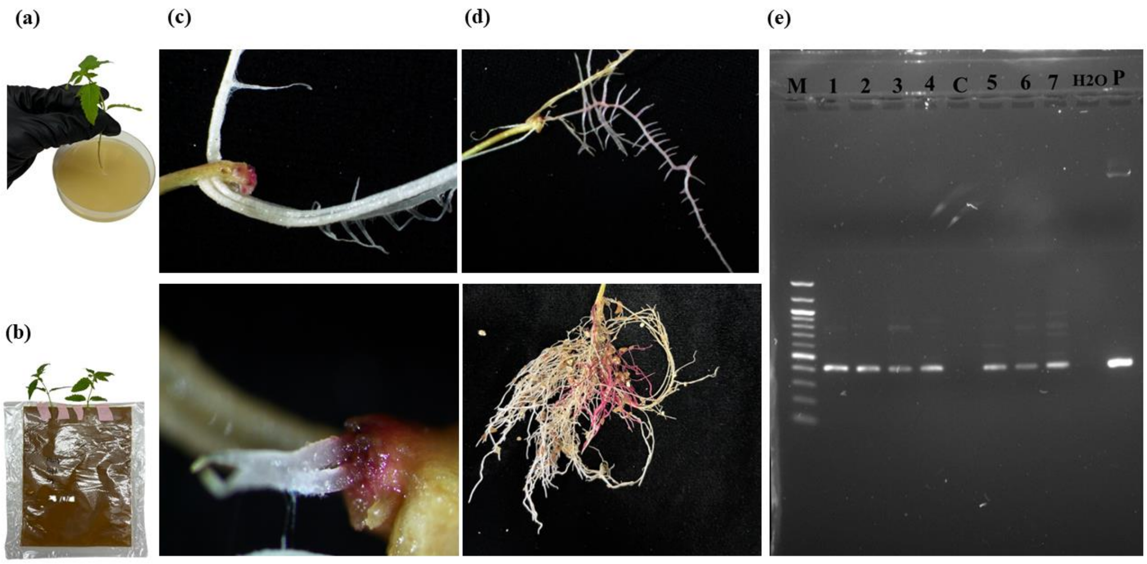


**Figure S1.** (a) Cannabis seedling infected with the specific bacteria and (b) placed in paper bags. (c) First sign of red (transformed) root. (d) A fully grown transformed hairy roots. (e) PCR results to validate transformation. M stands for a 100 bp DNA marker; 1–7 denotes red transgenic hairy roots; P is the 35S_Ω_RUBY plasmid used as a positive control; C stands for white hairy roots used as a control sample; and $H_2O$ is a negative control containing no template.
